# Pantoea in agriculture: Balancing promise and pathogenicity

**DOI:** 10.64898/2025.12.18.695097

**Authors:** Darin Holman, Enriquay Smith, Adeola Raji, Ameerah Abrahams, Mohamed-Deen Hendricks, Gerhard Basson, Augustine Daniel, Mbukeni Nkomo, Lee-Ann Niekerk, Morné Du Plessis, Arun Gokul, Ashwil Klein, Marshall Keyster

## Abstract

The genus Pantoea has gained attention as a promising biofertilizer and biocontrol agent, with strains like *Pantoea agglomerans* and *Pantoea dispersa* enhancing plant growth and suppressing pathogens. However, the same genus includes pathogenic species such as *Pantoea stewartii* (causing Stewart’s wilt in maize) and *Pantoea ananatis* (inducing center rot in onions). This duality raises critical concerns about the unregulated adoption of Pantoea in agriculture.

## The paradox of Pantoea in sustainable agriculture

The growing demand for sustainable agricultural practices has intensified interest in plant-associated microbes as alternatives to chemical fertilizers and pesticides [1]. Among these, the genus Pantoea has emerged as a particularly intriguing candidate, with demonstrated potential as both a biofertilizer and biocontrol agent. Certain species, such as *Pantoea agglomerans*, can fix nitrogen, solubilize phosphate, and suppress phytopathogens through antibiotic production [2]. However, this genus presents a striking paradox: while some strains promote plant growth, others are devastating pathogens responsible for diseases like Stewart’s wilt in maize (*Pantoea stewartii*) and center rot in onions (*Pantoea ananatis*) [3]. This duality raises critical questions about the safety and oversight of Pantoea-based agricultural products. Our literature analysis reveals significant geographical biases in research focus, with studies concentrated in developed regions while tropical agricultural zones remain understudied (Figure 1). Recent studies reveal that the genetic boundaries between beneficial and pathogenic strains are often blurred, with virulence factors such as type III secretion systems and phytotoxins appearing in otherwise “beneficial” isolates [4]. Compounding this risk, horizontal gene transfer (HGT) events can rapidly convert commensal strains into plant pathogens [5]. Despite these concerns, commercial applications of Pantoea are advancing with limited assessment of host range specificity or environmental persistence. The urgent need for a more rigorous, genomics-informed approach to Pantoea deployment is evident. As we advocate for microbial solutions to enhance food security, we must equally prioritize risk assessment to prevent unintended ecological consequences. This Comment highlights the dual nature of Pantoea, examines current gaps in biocontrol safety evaluation, and proposes actionable strategies to balance agricultural innovation with biosafety.

**Figure 1.**
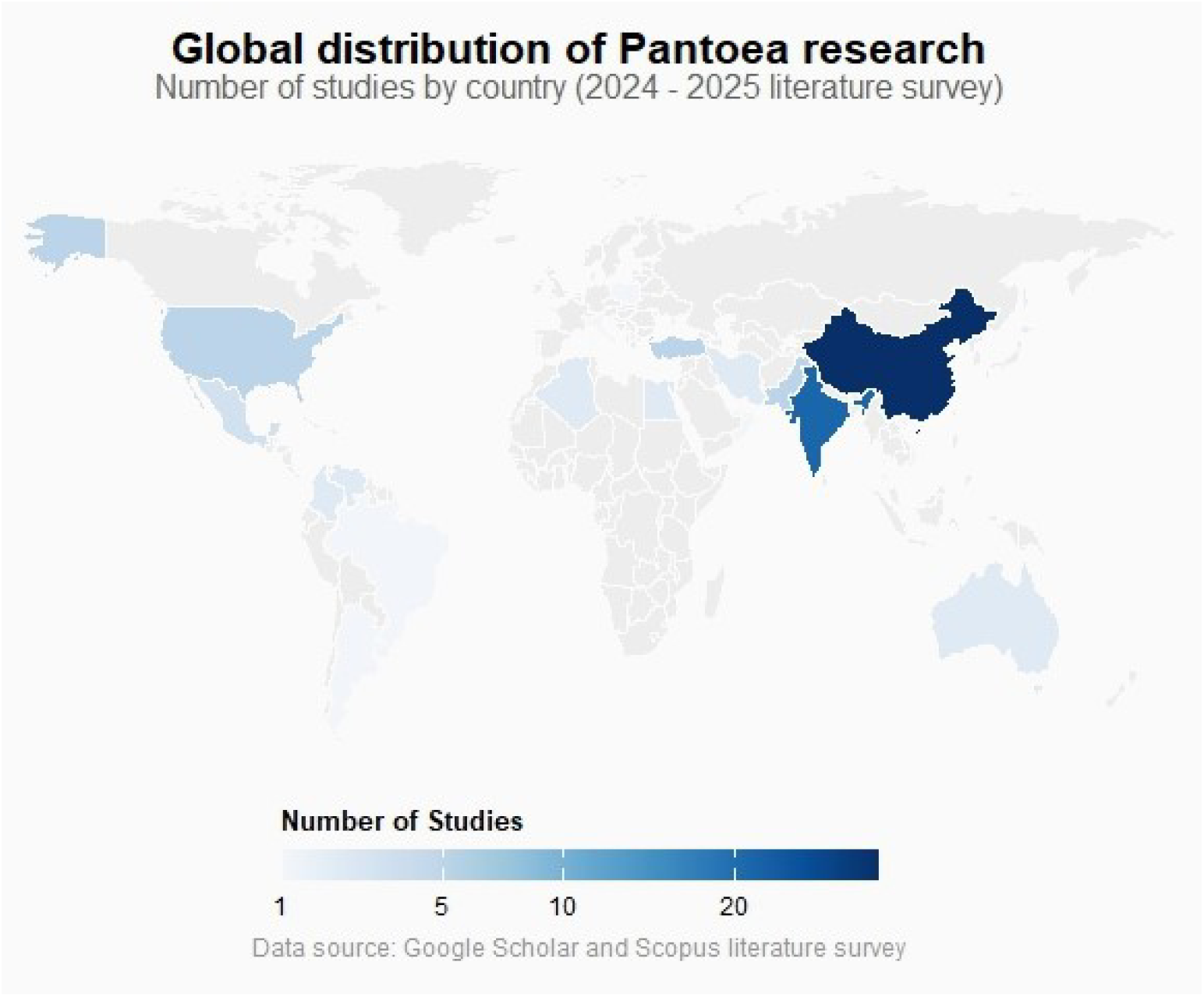
Global geographical distribution of Pantoea research, demonstrating significant regional biases in study locations. Research efforts are concentrated in North America, Europe, and Asia, with substantial gaps in tropical and developing regions where agricultural applications would be most relevant. All data were compiled from literature published between January 2024 and 10 October 2025 in Google Scholar and ScienceDirect. Statistical analyses and visualizations were performed using R version 4.5.1.

### Strain-specific risks demand rigorous assessment

The genomic plasticity of Pantoea species presents both opportunities and challenges for agricultural applications. While some strains exhibit remarkable plant growth-promoting traits, others harbor virulence determinants that pose significant risks to crop health. Comparative genomic studies have revealed that pathogenic and beneficial strains often differ by only a handful of key genes, including those encoding type III secretion systems (T3SS), phytotoxins, and effector proteins [4]. For example, *P. agglomerans* strain Eh318 produces the antibiotic pantocin A, which suppresses fire blight in apples, while closely related strains of the same species cause blight in rice and wheat. HGT further complicates risk assessment, as plasmids and genomic islands can rapidly disseminate virulence factors among strains. The pPATH plasmid family, commonly found in Pantoea spp., carries genes for host-specific virulence, including the hrp gene cluster essential for pathogenicity in *P. stewartia* [6]. Alarmingly, antibiotic resistance genes are frequently co-localized with these virulence factors on mobile genetic elements, raising concerns about the potential for environmental persistence and spread. Recent metagenomic surveys have detected Pantoea-associated virulence genes in agricultural soils, suggesting that introduced strains could exchange genetic material with indigenous microbiota. These findings underscore the need for comprehensive genomic screening of candidate strains prior to field application, particularly given the broad host ranges observed across Pantoea species (Figure 2).

**Figure 2.**
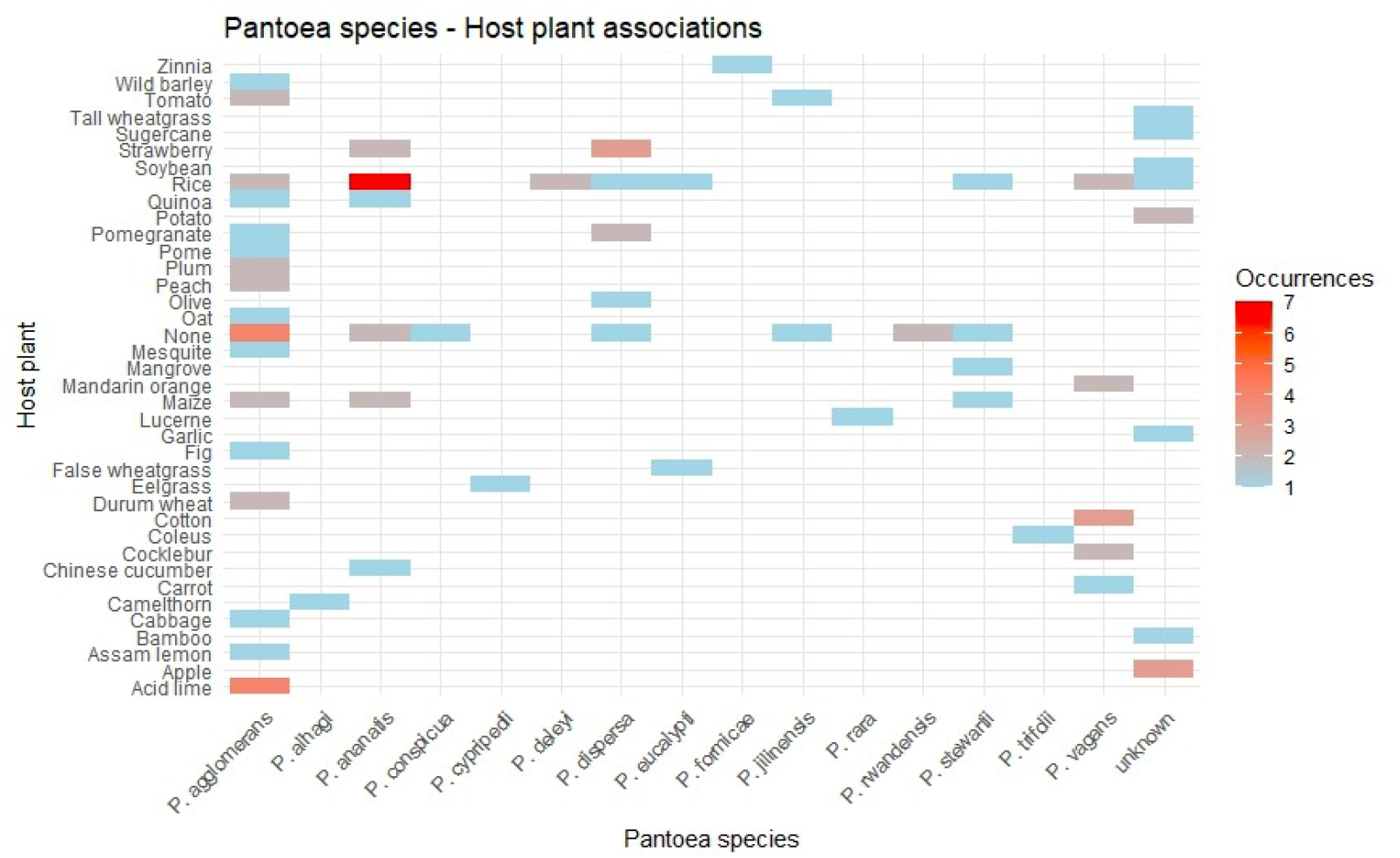
Host plant associations of Pantoea species, demonstrate broad host ranges and a lack of specificity in plant-microbe interactions. While species like *P. agglomerans* associate with numerous host plants, the vast majority of lesser-studied species also show promiscuous associations across diverse plant families. This non-specificity underscores the potential for unintended ecological consequences when deploying Pantoea strains as biofertilizers or biocontrol agents.

### The need for comprehensive testing before commercialization

Current approaches to evaluating Pantoea-based biofertilizers often suffer from narrow experimental designs that fail to assess potential off-target effects. Most studies focus on a single crop system under controlled conditions, neglecting the complex interactions that occur in diverse agricultural ecosystems. For instance, *P. ananatis* strain JCC14, showed growth promotion effects in maize but caused rot symptoms in onion^7^. This host-dependent variability necessitates systematic testing across multiple plant families under field-realistic conditions, especially concerning given the context-dependent effects observed even on original host plants (Figure 3). Emerging technologies offer promising solutions for improved risk assessment. Whole genome sequencing coupled with machine learning algorithms can identify high-risk genomic signatures predictive of pathogenicity in Pantoea [8]. For example, comparative pan-genome analyses have identified accessory genome components associated with virulence in Pantoea [7]. High-throughput phenotyping platforms could screen candidate strains against hundreds of plant varieties simultaneously, while microbiome interaction studies would reveal potential impacts on soil microbial communities^9^. Such comprehensive approaches should become standard practice before regulatory approval, particularly important given the dual functionality many species exhibit under stress conditions (Figure 4).

**Figure 3.**
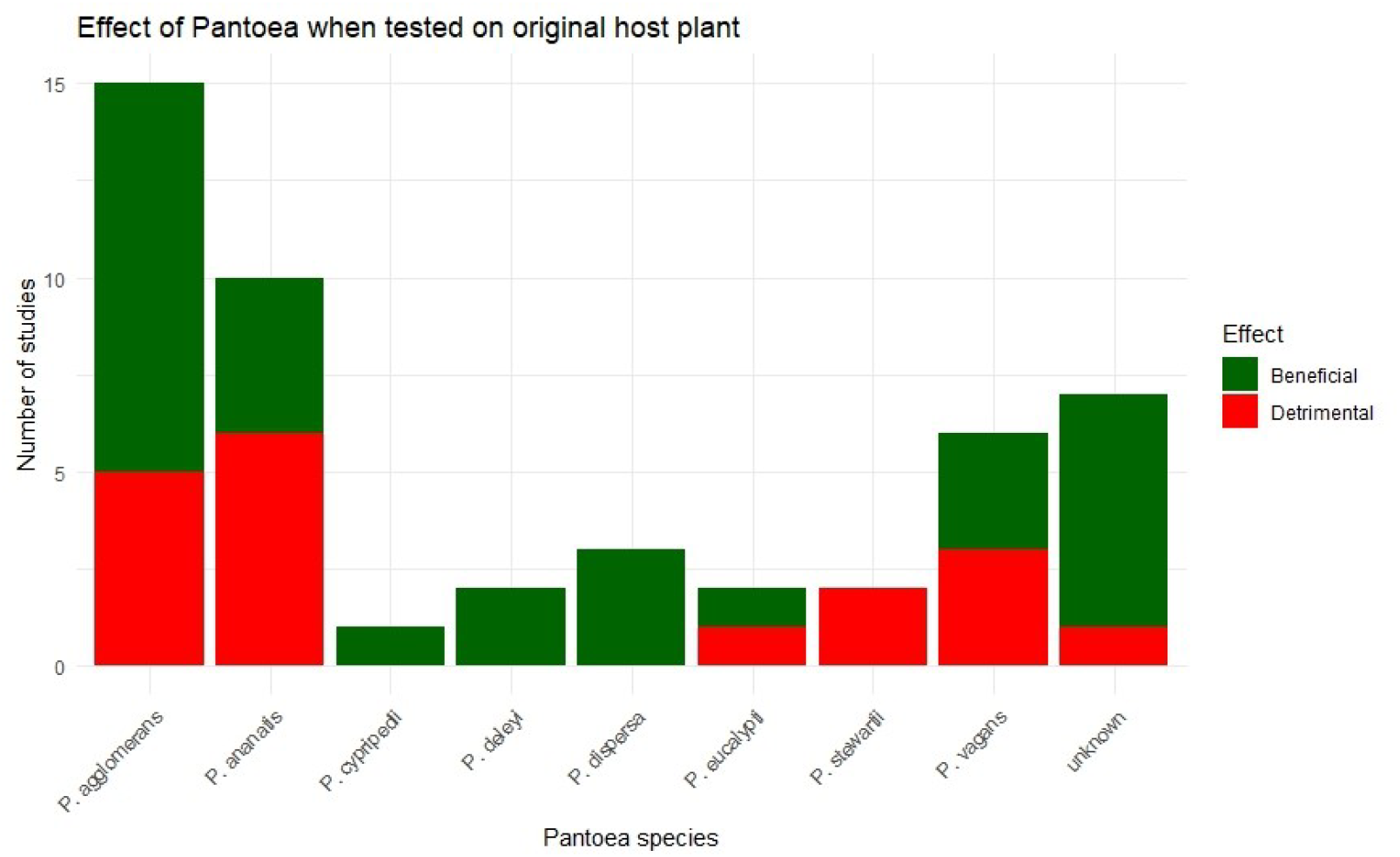
Context-dependent effects of Pantoea species when tested on their original host plants. Strikingly, many species exhibited both beneficial and detrimental effects even on the plants from which they were originally isolated. Only a minority of species demonstrated exclusively beneficial effects, while a single species was exclusively detrimental. This variability underscores the strain-specific and condition-dependent nature of Pantoea-plant interactions, challenging simplistic categorization of species as purely beneficial or pathogenic.

**Figure 4.**
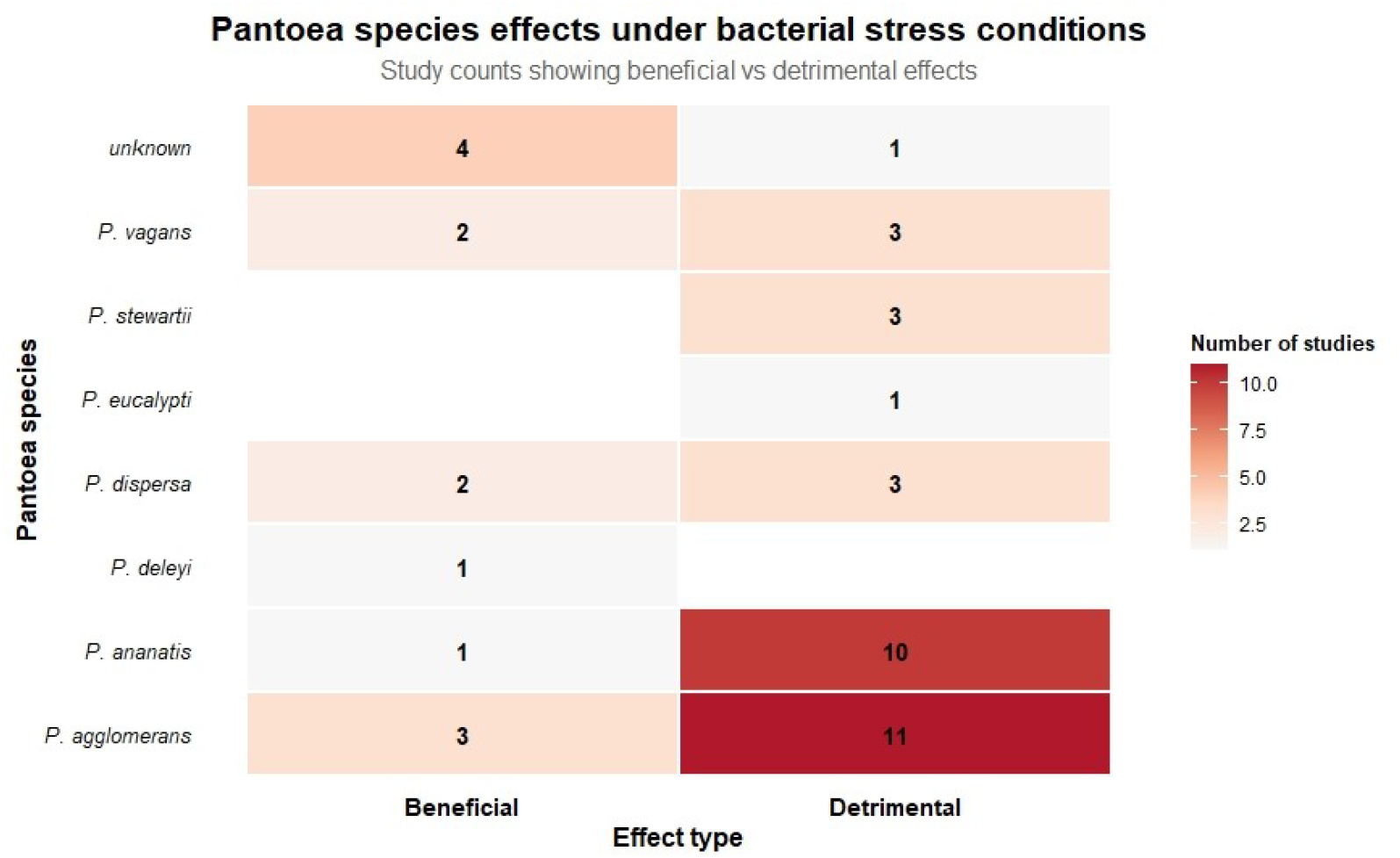
Effects of Pantoea species under bacterial stress conditions, revealing alarming functional duality. The heatmap demonstrates that multiple species exhibit both beneficial and detrimental effects depending on context, with only one species displaying exclusively beneficial effects and two species showing exclusively detrimental effects under stress conditions. This dual functionality in stress response highlights the unpredictable nature of Pantoea species and underscores the potential risks of deploying these bacteria without comprehensive stress testing.

### A call for stricter regulation and interdisciplinary collaboration

The current regulatory landscape for microbial inoculants remains fragmented, with significant gaps in safety assessment protocols. Unlike chemical pesticides, microbial products often receive approval based on limited toxicity data without adequate evaluation of ecological impacts [10]. A harmonized international framework is urgently needed, incorporating: (1) mandatory whole genome sequencing and annotation of commercial strains, (2) multiseason field trials across diverse agroecosystems, (3) assessment of HGT potential through conjugation experiments, and (4) monitoring programs for post-release environmental impacts. The establishment of a global Pantoea registry, integrating genomic and phenotypic data with geographical distribution information, would significantly improve risk assessment capabilities. This effort requires collaboration between microbiologists, computational biologists, plant pathologists, and regulatory agencies. Industry partners must commit to transparent data sharing, while funding agencies should prioritize research on microbial risk assessment methodologies. The analysis presented here underscores that without such coordinated efforts, the potential benefits of Pantoea in sustainable agriculture may be outweighed by unintended ecological consequences.

## Conclusion

The promise of Pantoea in sustainable agriculture must be balanced against its demonstrated capacity for plant pathogenesis. While certain strains offer genuine benefits, the genus’ genomic fluidity and capacity for rapid evolution demand cautious, science-based approaches to commercialization. Three critical steps must precede widespread adoption: implementation of robust genomic screening protocols, development of host-range prediction models, and establishment of international safety standards. Only through such rigorous measures can we harness Pantoea’s potential while safeguarding global food security and ecosystem health. The time to address these challenges is now, before uncontrolled applications lead to irreversible agricultural consequences.

## Funding

MK received grant funding from the DSI-NRF Centre of Excellence in Food Security (Grant ID: 25202). MK also received research grant funding from the National Research Foundation of South Africa (Grant number: 151752). AK and MK received joint-funding from the Department of Science and Innovation-Technology Innovation Agency (Grant number: GB0200090). DH, ES, and AR were funded by the Mastercard Scholars Foundation of the University of the Western Cape. MH received PhD funding from the National Research Foundation of South Africa. GB was funded by a Postdoctoral fellowship of the National Research Foundation of South Africa. AD was funded by the UWC Research Chair in Sustainable Agriculture (Grant number: GB0200088). The funders had no role in study design, data interpretation, or publication decisions.

### SDGs

SDG2, SDG9, SDG12, SDG15, and SDG17

## Acknowledgements

The authors gratefully acknowledge the organizers and participants of the 1st International conference on Pantoea (Skukuza, South Africa, 2025) for their insightful discussions that helped shape the perspectives presented in this commentary. The conference served as an invaluable platform for exchanging knowledge about this complex genus and its agricultural implications. Furthermore, the authors would like to thank the University of the Western Cape and the University of Free State for infrastructure and administrative support.

## Data availability

No new data were generated or analyzed in this study. All information discussed in this article is derived from previously published sources, which are cited within the manuscript.

